## Supplementary figures and images for "A comprehensive two-hybrid analysis to explore the *L. pneumophila* effector-effector interactome"

### Figure EV1

#### Figure EV1 - iBFG-Y2H barcode representation and correlation of fusion barcode tags

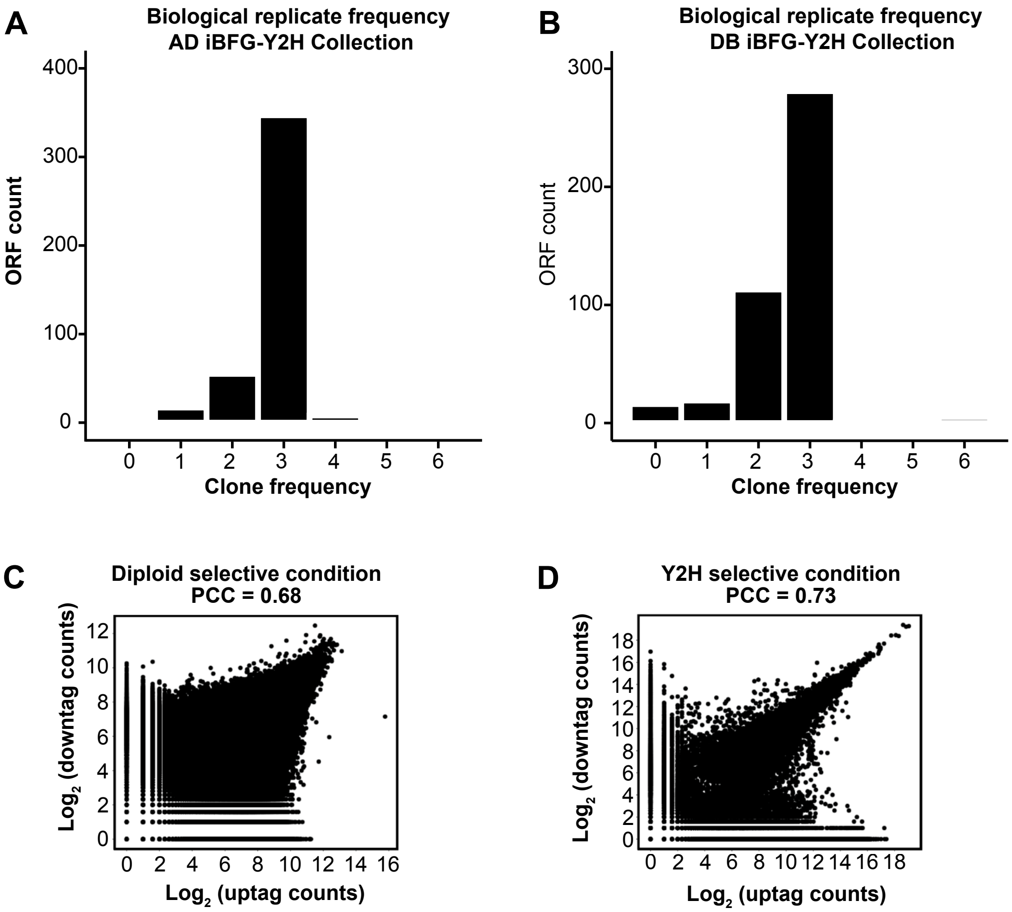

### Figure EV2

##### Figure EV2 - Retest of iBFG-Y2H interactions with *L. pneumophila* effectors

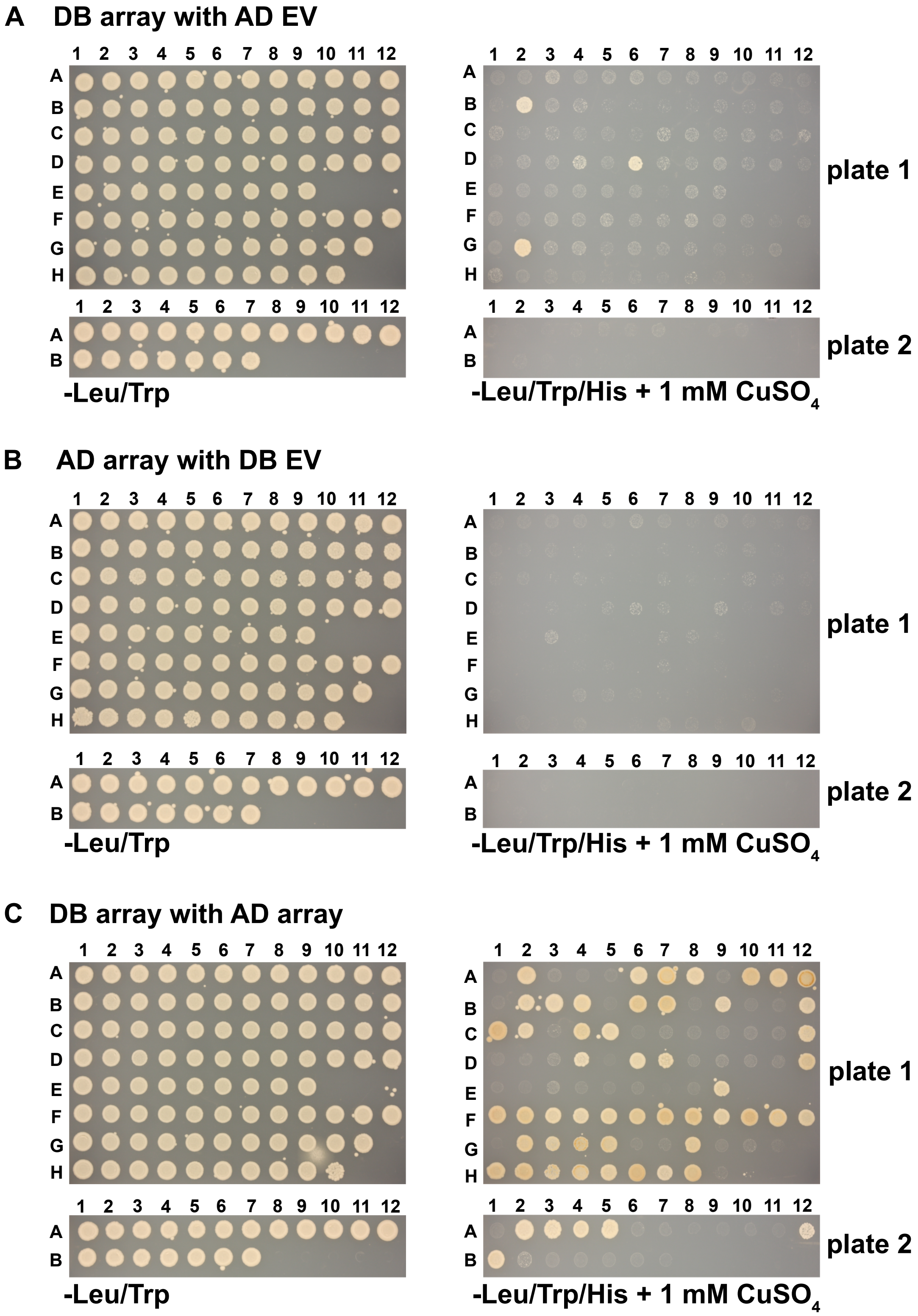
